# The transcription factor Capicua maintains the oocyte polarity in the panoistic ovary of the German cockroach

**DOI:** 10.1101/2024.10.21.619414

**Authors:** Nashwa Elshaer, Jorge Escudero, Maria-Dolors Piulachs

## Abstract

The establishment of the symmetry axis is crucial for the development of all organisms. In insects, this process begins early in oogenesis with the correct distribution of the mRNAs and proteins in the oocyte. One protein that plays a role in organizing this distribution is the transcription factor Capicua (Cic). Cic has been studied in the context of oogenesis and embryonic development in *Drosophila melanogaster*. It is maternally expressed, begins essential for establishing the dorsoventral axis, and functions as a transcriptional repressor. Although the Cic sequences are conserved across species, their function in other types of insect ovaries is still little known. We wondered whether the function of Cic in insects has been maintained through evolution despite the ovary type or if it has been modified in parallel to the ovary evolution. To address this, we studied the Cic function in a phylogenetically basal insect, the cockroach *Blattella germanica*, a species with panoistic ovaries. Our findings show that *B. germanica* Cic is essential for oocyte development and the maturation of ovarian follicles. A loss of Cic function leads to disrupted cytoskeletal organization, defects in anterior-posterior polarity, and compromised follicle integrity. The conservation and functional divergence of Cic across different species suggest evolutionary adaptations in the mechanisms of insect oogenesis.

## 1. Introduction

Oogenesis is a crucial process in the animal kingdom, ensuring species continuity. Therefore, it must be regulated with great precision. A good model for studying oogenesis is the insect ovary, especially, the panoistic ovary, characterized by the absence of nurse cells escorting the germinal cell or oocyte. Instead of this organization, each ovarian follicle only consists of an oocyte surrounded by a monolayered follicular epithelium (Büning, 1994). The panoistic ovary is characteristic of phylogenetically basal insects like the cockroach *Blattella germanica*, our insect model (Belles et al., 2024), and it is the most common ovarian type among invertebrates and vertebrates (Telfer, 1975). A distinguishing feature of *B. germanica* oogenesis is that the basal ovarian follicle (BOF), from each ovariole, is the only one that matures in each reproductive cycle, while the other ovarian follicles remain in the vitellarium, awaiting their chance to occupy the basal position in subsequent cycles (Belles et al., 2024; Irles & Piulachs, 2014; Tanaka & Piulachs, 2012) (Fig. 1). The BOFs begin to grow during the last nymphal instar. When vitellogenesis starts in 3-day-old adults, the BOF grows exponentially (Fig. 1), increasing its size tenfold. During this process, the follicular cells in the BOF proliferate, encompassing the oocyte growth. Coinciding with vitellogenesis, the follicular cells arrest cytokinesis, becoming binucleate and entering into endoreplication. By the end of the cycle, there is an amplification of the chorion genes, which are synthesized in a few hours to complete the egg (Fig. 1).

**Figure 1.**
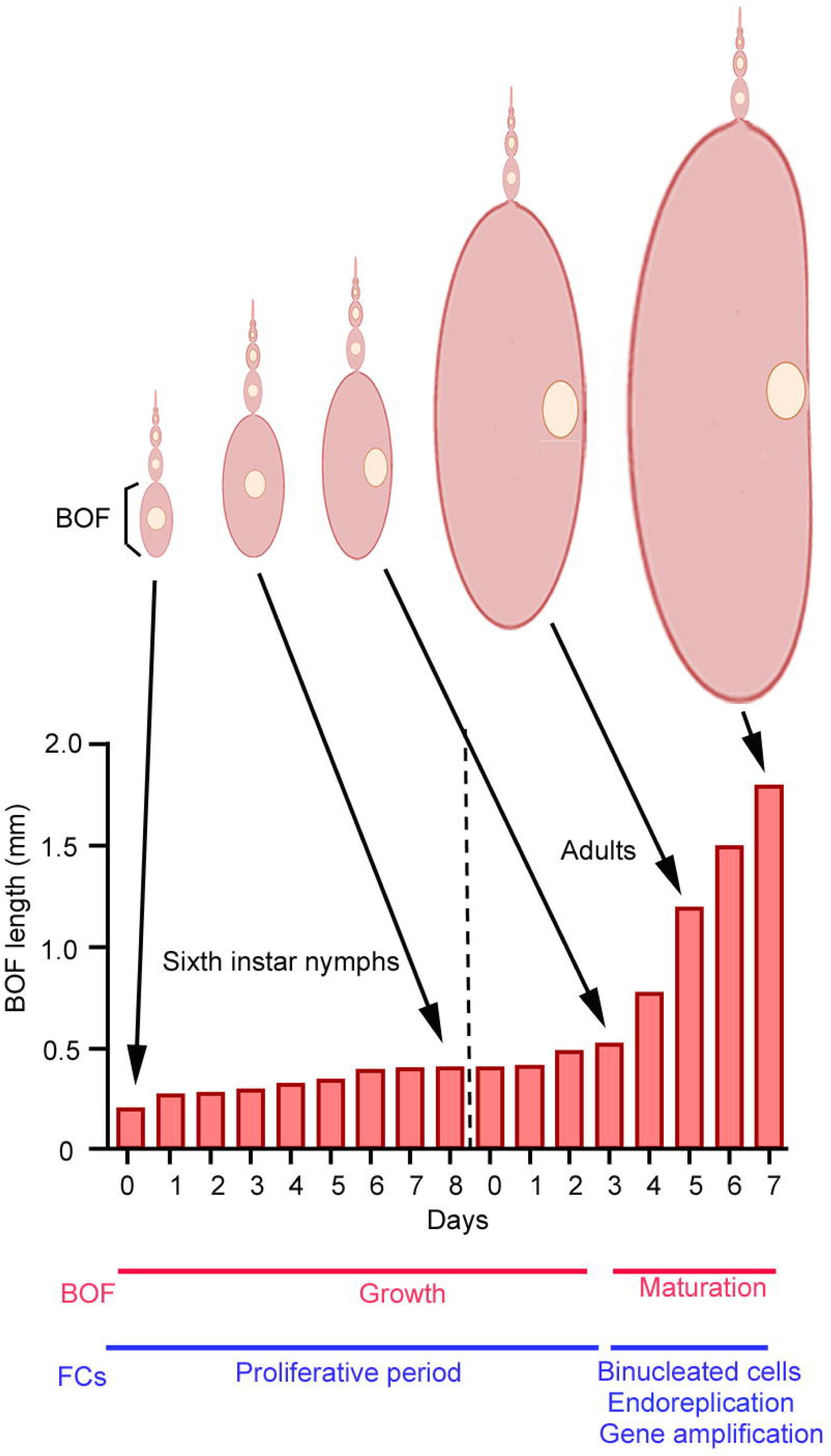
Changes in the *Blattella germanica* basal ovarian follicle, during the gonadotropic cycle. Geometric growth of the basal ovarian follicle (BOF), in key moments of the gonadotropic cycle. The BOFs grow linearly during the last instar nymphs, until 3-day-old adults. During this period, the follicular cells (FCs) surrounding the basal oocyte proliferate. In 3-day-old females, the vitellogenesis begins, and the BOF grows exponentially, increasing ten times its size. The oocyte nucleus migrates to the ventral pole, where the germinal band will be formed. The FCs arrest cytokinesis becoming binucleated, and enter endoreplication. In the last days of the cycle, these cells amplify some genome regions to improve the synthesis of the chorion proteins.

During oocyte growth, the organization of the future embryo is determined. The positioning of the nucleus and cell organelles, as well as the localization of specific proteins and RNAs within the oocyte, determine the establishment of oocyte polarity. This polarity will be essential later, during the embryogenesis, to reach a correct body plan development (Bashirullah et al., 1998; Bradley et al., 2000; Wilson et al., 2011). *Drosophila melanogaster*, with meroistic polytrophic ovaries, is the species in which oocyte polarity has been most thoroughly studied. The ovarian follicles in a polytrophic ovary are asymmetric, as they are formed by a cluster of nurse cells and the oocyte, all of them, surrounded by a monolayer of follicular cells (Büning, 1994). In this species, oocyte asymmetry is established in mid-oogenesis by the cell-cell interactions between the oocyte and follicular cells, which ultimately determine the final position of the oocyte nucleus. The displacement of the oocyte nucleus to a dorsal-anterior position is facilitated by the distribution of microtubules in the ooplasm, ensuring the proper localization of mRNAs, like *Oskar* or *bicoid,* in the ooplasm (Roth & Lynch, 2012; Wilson et al., 2011). Conversely, in panoistic ovaries, like those in *B. germanica*, the ovarian follicles exhibit a bilateral symmetry. The changes in symmetry that occur during oocyte development are subtle and occurs in adults, and information about this process remains limited. In *Acheta domesticus* (Netzel 1968; Bradley et al., 2000) and *B. germanica* (Tanaka 1973; Zhang & Kunkel, 1992), changes in the position of the oocyte nucleus, facilitate distinguishing the dorsoventral axis. In both species, the nucleus moves from a central to the ventral position. In *B. germanica*, the oocyte nucleus is maintained in a mid-ventral position (Tanaka, 1973). In addition, the asymmetric distribution of F-actins in the oocyte membrane, together with the maintenance of patency in the ventral follicular cells, facilitate the identification of the ventral side of the BOF, when the oocyte reaches its final size (Zhang & Kunkel, 1992).

The formation of the anteroposterior axis in oocytes of panoistic ovaries is even less well known. One of the first indications of the presence of an anteroposterior axis was given by Neztle (1968) in *A. domesticus* when he described the displacement of the nucleus towards a ventral, slightly posterior position. Later, Bradlez and collaborators (Bradlez et al., 2000), also in *A. domesticus*, described the formation of Balbani bodies in both poles, anterior and posterior. Initially in the anterior pole and later in the posterior, this determines the special localization of the RNAs and proteins. In the case of *B. germanica*, we described a population of follicular cells in the apical pole of the BOF, which differentiate at the end of the cycle to synthesize the egg structures (Pascual et al., 1992; Irles et al., 2009). This limited available information requires further investigation about the organization of oocytes and eggs in insects with panoistic ovaries.

We have adopted the panoistic ovary of *B. germanica* as a model, to study the function of the transcription factor Capicua (Cic), which acts as a repressor and is structurally conserved from cnidarians to vertebrates (Jiménez et al., 2000; Lam et al., 2006; Rodrıguez-Muñoz et al., 2022). In *D. melanogaster*, two Cic isoforms have been described. One short, Ci*c*-S, considered the canonical isoform (Rodrıguez-Muñoz et al., 2022) reported as a maternally expressed gene that is required for the establishment of dorsoventral patterning of the eggshell and the embryonic termini during embryogenesis (Goff et al., 2001; Jiménez et al., 2000), the other longer isoform Cic-L, the most common between metazoan, is expressed in all tissues (Rodrıguez-Muñoz et al., 2022). In *Tribolium castaneum*, with meroistic telotrophic ovaries, the function of Cic in the terminal patterning is conserved, although its depletion does not affect the dorsoventral axis (Prido lh. et al., 2017)

Our findings indicate that Cic is necessary for maintaining polarity in the ovarian follicles of *B. germanica* and for stabilizing the distribution of F-actin in oocytes, a function that seems to be performed early during oocyte development.

## 2. Materials and Methods

### 2.1. Insects

*Blattella germanica* (L.) specimens were obtained from a colony reared in the dark at 29 ± 1°C and 60–70% relative humidity (Bellés et al., 1987). Adult females were maintained with males during all the first gonadotrophic cycle, and mating was confirmed at the end of the experiments by assessing the presence of spermatozoa in the spermathecae. All dissections and tissue sampling were carried out on carbon dioxide-anesthetized specimens. Tissues were frozen on liquid nitrogen and stored at -80°C until use.

### 2.2. Cloning and Sequencing

A fragment of *B. germanica capicua* (*cic*) sequence (493 bp) was obtained from an mRNA library available in the laboratory. The sequence of *cic* was completed by 3’- and 5’- rapid amplification of cDNA ends (RACE), according to the kit protocol (Ambion, Huntingdon, Cambridgeshire, UK). The amplified fragments were cloned into the pSTBlue-1 vector (Novagen, Madison, WI, USA) and sequenced by Sanger (accession number in NCBI: LN623700).

### 2.3. Sequence comparisons and phylogenetic analysis

Sequences of insect Cic protein, used in the phylogenetic analysis, were obtained from GenBank. The search was enlarged by Blast to NCBI, using the *B. germanica* and *D. melanogaster* Cic as queries. Cic sequences were selected taking representatives of diverse insect taxa. Priority was given to species with well-annotated genomes to ensure high-quality data. The accession numbers of the sequences used are listed in Supplementary Table 1. The amino acid sequences were aligned using MAFFT (Katoh et al., 2019). The resulting alignment was analyzed by the PHYML 3.0 program (Guindon et al., 2010) based on the maximum-likelihood principle, using a JTT matrix, four substitution rate categories with a gamma shape parameter of 1.444, and using empirical base frequencies and estimating proportions. The data was bootstrapped for 100 replicates.

The domains NLS, TUDOR, N1, HMG, C2, and C1 in the Cic sequences of the studied species were localized based on the description done in *H. sapiens* and *D. melanogaster* in Webb et al, 2022, and Rodriguez-Muñoz et al., 2022 (See Fig. S1).

### 2.4. RNA extraction and expression studies

Total RNA was isolated using the GenElute Mammalian Total RNA kit (Sigma, Madrid, Spain). An amount of 400 ng from each RNA extraction was DNAse treated (Promega, Madison, WI, USA) and reverse transcribed with Superscript II reverse transcriptase (Invitrogen, Carlsbad, CA, USA) and random hexamers (Promega). RNA quantity and quality were estimated by spectrophotometric absorption at 260/280 nm in a Nanodrop Spectrophotometer ND-1000® (NanoDrop Technologies, Wilmington, DE, USA).

Expression of *cic* was determined by quantitative real-time PCR (qRT-PCR) in the last nymphal instar and the adult stage during the first gonadotrophic cycle. PCR primers used in qRT-PCR expression studies were designed using the Primer Express 2.0 software (Applied Biosystems, Foster City, CA, USA), and are detailed in Supplementary Table 2. The *B. germanica actin-5c* mRNA (Accession number AJ862721), was used as a reference. qRT-PCR reactions were performed using the SYBR Green Supermix (BioRad) containing 200 nM of each specific primer. Amplification reactions were carried out at 95°C for 2 min, and 40 cycles of 95°C for 15 s and 60°C for 30s, using MyIQ Single Color RTPCR Detection System (BioRad). After the amplification phase, a dissociation curve was carried out to ensure that there was only one product. Levels of mRNA were calculated relative to the respective reference gene, using the 2^-ΔΔCt^ method and 2^-ΔCt^ method (Irles et al., 2009; Livak & Schmittgen, 2001; Maestro et al., 2005). Results are given as mRNA copies per 1000 copies of *actin-5c* mRNA and correspond to three biological replicates.

### 2.5. RNA interference

To deplete *cic* mRNA, and asses the specificity of the phenotype, two different dsRNAs (ds*cic*-1 and ds*cic*-2) were designed and synthesized as previously described (Ciudad et al., 2006). The ds*cic*-1 was 324 bp in length (from nucleotides 5,874 to 6,197) and the second one, ds*cic*-2, 392 bp (from nucleotides 4,366 to 4,757). A dsRNA, corresponding to 307 bp of the *Autographa californica* nucleopolyhedrovirus sequence (GenBank: K01149), was used as a control (dsMock). The corresponding cDNAs were amplified by PCR and cloned into the pSTBlue-1 vector. Newly-emerged sixth instar nymph females, or newly-emerged adult females, were independently treated with 1 μg of ds*cic*-1 or 1 μg ds*cic*-2. Since the same ovary phenotype was observed regardless of which dsRNA was used, we will refer to RNAi treatments collectively as ds*cic*. The same dose and conditions applied in ds*cic* treatments were applied to dsMock-treated females.

### 2.6. Immunohistochemistry

After dissection, ovaries (n = 4, for each experiment and treatment) were fixed immediately for 2 h with 4% paraformaldehyde in PBS, washed in PBT (PBS; 0.3% Triton-X100), treated with 50 µg/ml proteinase K for 2 min, washed for 2 min in 2 mg/mL glycine in PBT, washed for 10 min in PBT and fixed again for 20 min in the same solution. Then the ovaries were washed three times for 10 min with PBT (PBS; 0.1% Triton-X100, 0.1% BSA). After three washes with PBT the ovaries were saturated for 1 h at room temperature in PBTBN (PBS; 0.1% Triton-X100, 0.5% BSA and 5% normal goat serum), and incubated overnight at 4°C with the primary antibody. The primary antibodies used were rat anti-Cic (1:200, generated against the C-terminal half of *D. melanogaster* Cic protein, kindly provided by Jordi Casanova; Molecular Biology Institute of Barcelona, CSIC), mouse anti-EGFR (15µg/mL) and mouse anti-Eya (1:50). Tissues were washed with PBTBN three times and incubated for 2 h with Alexa-Fluor 647 goat anti-mouse or anti-rat IgG (Molecular Probes, Carlsbad, CA, USA) diluted 1:400 in PBTBN. For F-actin visualization, ovaries were incubated for 20 min in 1µg/mL phalloidin-TRITC (Sigma), and after washing they were mounted in UltraCruz™ Mounting Medium (Santa Cruz Biotechnology^®^, inc., Delaware CA, USA), which contains DAPI for DNA staining. Samples were observed by epifluorescence microscopy using a Zeiss AxioImager.Z1 microscope (ApoTome), using the software Zen 2012, blue edition (Carl Zeiss MicroImaging). The relative fluorescence intensities of the Cic labeling, DAPI, and phalloidin-TRITC, were graphically quantified using the *Zen 2.5D analysis tool* (Carl Zeiss Microscopy).

### 2.7. Statistics

Data are expressed as mean ± standard error of the mean (S.E.M.). Statistical analysis of gene expression values was carried out using the REST 2008 program (Relative Expression Software Tool V 2.0.7; Corbett Research) (Pfaffl et al., 2002). This program makes no assumptions about the distributions, evaluating the significance of the derived results by Pair-Wise Fixed Reallocation Randomization Test tool in REST (Pfaffl et al., 2002). The sample group was considered significantly different from the control group when P(H1) = 0.000.

## Results

### 3.1. *Blattella germanica* has a structurally conserved Capicua ortholog

*capicua* (*cic*) cDNA from *B. germanica*, was amplified, cloned, and sequenced from ovarian tissues. The complete sequence of *B. germanica cic* has 7,631 bp with an ORF encoding for a protein of 2,279 amino acids (nucleotide positions 553-7,392). It is similar to the Cic-L isoform from *D. melanogaster*, and similar to the sequences of other insect species available in databanks (Fig. S1). From the sequence comparisons, the Dipteran Cic included in the analysis, could be an exception since they have a sequence corresponding to the short isoform. *B. germanica* Cic possesses the HMG-box domain (from amino acid 1,206 to 1,274) characteristic of Cic proteins highly conserved between insect species (Fig. S1). In addition, the NLS, TUDOR, N1 and N2 domains, the helical C1 module, and the C2 MAPK docking site, which helps DNA binding (Webb et al., 2022), are also present in the insect species analyzed, showing also a high degree of conservation (Fig. S1).

Maximum-likelihood analysis of Cic sequences from insects available in the literature and databases gave a phylogenetic tree (Fig. 2) with two clades, clearly separated corresponding to hemimetabolous (purple) and holometabolous (blue) insect species. The topology of the phylogenetic tree of Cic sequences overlaps with the currently accepted phylogeny of the included species (Misof et al., 2014). All insect orders appeared grouped with high consistency in the nodes. Interestingly, the Cic sequences from Lepidoptera and Diptera form two clusters with longer branches, indicating more extensive changes and, consequently, a faster divergence rate compared to the Cic proteins from other insect orders. The Cic-S and Cic-L from *D. melanogaster* appear to be grouped in the same node (Fig. 2), suggesting they share the same origin (Fig. 2).

**Figure 2.**
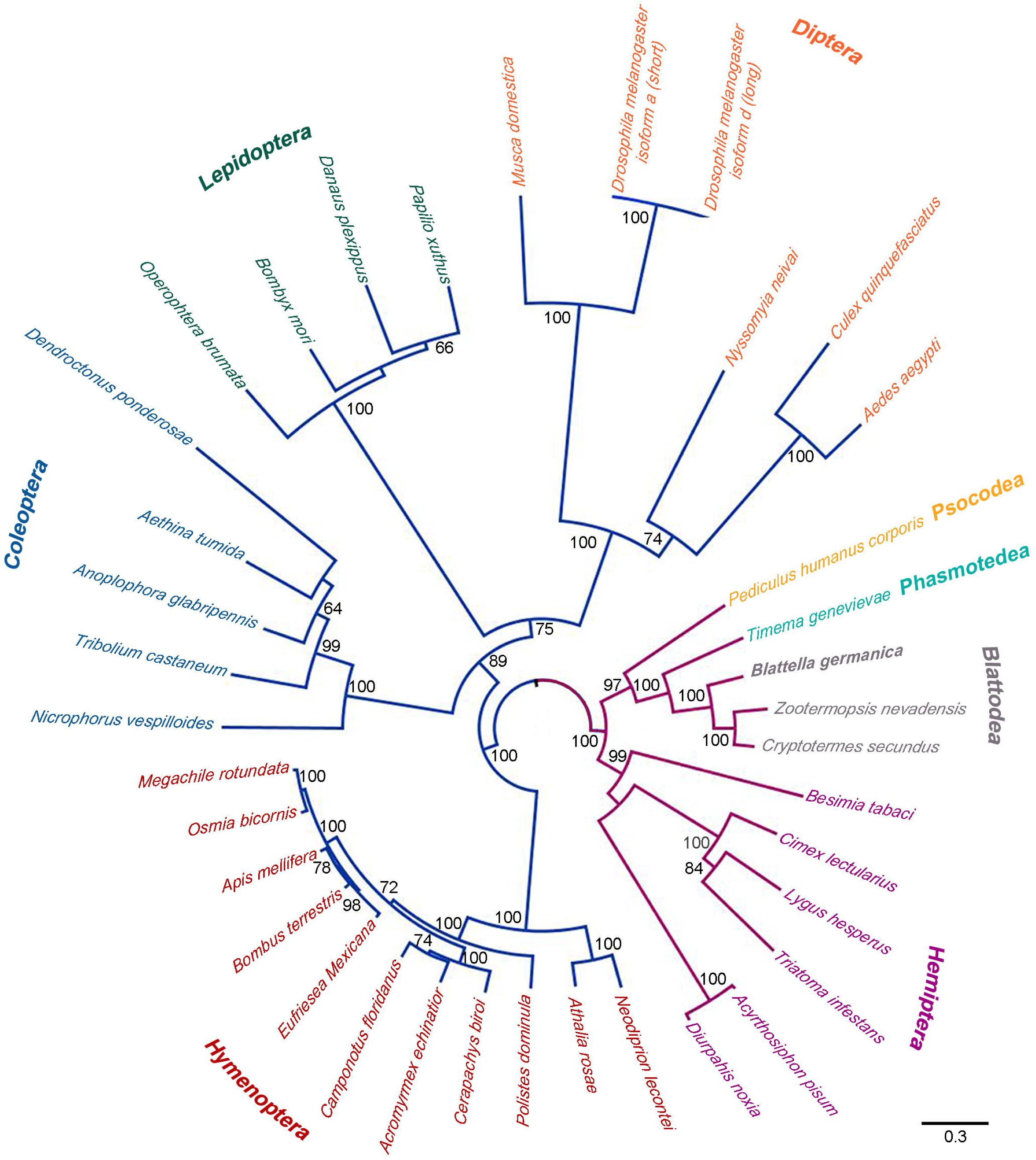
Phylogenetic analysis of the insect transcription factor Capicua, The analysis is based on the maximum-likelihood principle, with the amino acid substitution model, and was carried out using insect Cic sequences available in public databases. The Cic sequence of *B. germanica* is highlighted in bold grey. Blue branches indicate species belonging to holometabolous insects, while purple branches represent species belonging to hemimetabolous insects. Only bootstrap with values greater than 60 are shown. The complete species names and accession numbers of the sequences are listed in Table S1. The scale bar indicates the number of substitutions per site.

### 3.2. *cic* is highly expressed in the ovary during the last nymphal instar

The expression of *cic* mRNA was studied in different tissues from 3-day-old adult females, namely the ovary, thoracic ganglia, colleterial glands, fat body, brain, and midgut (Fig. 3A). *cic* is expressed in all tissues analyzed, with the highest levels in the brain and ovary. In ovaries, the expression pattern of *cic* does not show significant changes (Fig. 3B). In general, in ovaries from sixth instar nymphs, *cic* mRNA levels are higher compared to the levels of the last day of the fifth nymphal instar. After the imaginal molt, *cic* expression tends to decrease, reaching the lowest levels just before oviposition (Fig. 3B). These expression patterns suggest that the main function of Cic in the ovary occurs during the last nymphal instar.

**Figure 3.**
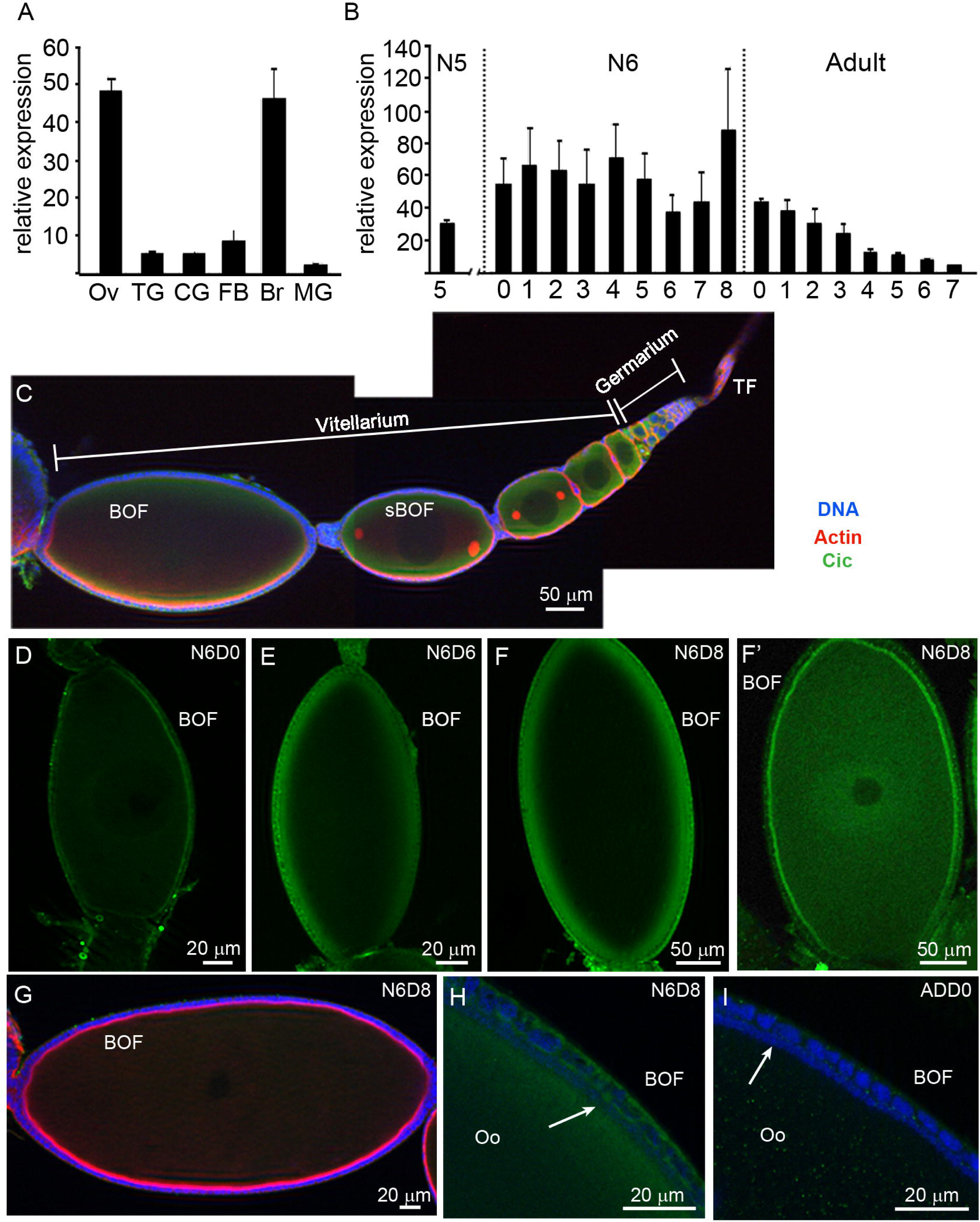
*B. germanica capicua* (*cic*). Expression and localization. (A) Relative expression of *cic* mRNA in ovary (Ov), thoracic ganglia (TG), colleterial glands (CG), fat body (FB), brain (Br), and midgut (MG) from adult females. (B) Relative expression of *cic* in the ovary during the sixth (last) instar nymph and in the adult. The expression on the last day of the fifth nymphal instar ovaries was also measured. Data in (A) and (B) are expressed as copies per 1000 copies of *actin-5c* (n = 3-4). (C) Immunolocalization of Cic in an ovariole from a 6-day-old sixth instar nymph showing the asymmetric distribution of the actin in the basal ovarian follicle (BOF) and sub-basal ovarian follicle (sBOF), TF: terminal filament. (D-F) Immunolocalization of Cic in basal follicles from 0-day-old (D), 6-day-old (E), and 8-day-old (F and F’) sixth instar nymphs, showing the changes of Cic distribution related to the age of the female, from an accumulation in the oocyte membrane (D, E) to be extended by the cytoplasm and nucleus (F’). In these images, the exposure time is the same, to allow comparisons. (G) Weakly Cic labeling in the follicular cells of the anterior and posterior poles in the BOF from 6-day-old sixth instar nymphs. (H-I) Optical section of BOFs from 8-day-old sixth instar nymph (H) and 0-day-old adult (I), showing differences in Cic distribution and labeling intensity within the ooplasm and the follicular cells. The arrows, in both images, indicate the layer of bacteriocyte endosymbionts. Oo: ooplasm. The anterior pole in C and G is towards the right, and in D-F it is to the top of the image. Cic: green, F-actin: red, DAPI: blue.

### 3.3. Cic protein localizes in somatic and germ cells

A heterologous antibody against the *D. melanogaster* Cic protein was utilized to localize Cic in *B. germanica* ovaries. Cic labeling was detected in ovaries of sixth instar nymphs, in all ovarian follicles throughout the ovariole, encompassing germinal and somatic cells (Fig. 3C). However, there are differences in the distribution of Cic in the growing oocyte. In the basal follicle of 0-day-old sixth instar nymphs, Cic labeling appeared as a thin layer adjacent to the oocyte membrane (Fig. 3D), with lower intensity at both the anterior and posterior poles of the oocyte. Later, in 6-day and early 8-day-old sixth instar nymphs (Fig. 3E and F, respectively), Cic labeling became more widespread through the ooplasm and accumulated as a thick layer at the oocyte boundary. This labeling is displayed in all ovarian follicles, from the basal to the most distal, as well as in the germarium (Fig. 3C). In 8-day-old sixth instar nymphs, as the molt to adult approaches, Cic labeling appears in the oocyte nucleus, spreading gradually with a faint signal throughout the cytoplasm, becoming more prominent at the oocyte membrane (Fig. 3F’). After the imaginal molt, Cic labeling in the oocyte becomes very weak (Fig. 3G). Concerning follicular cells, in 8-day-old sixth instar nymphs, Cic appears abundant in the cytoplasm (Fig. 3H). In contrast, in 0-day-old adults, the labeling in the follicular cells became fainter and is concentrated at the basal pole (Fig. 3I). Overall, these findings indicate that Cic localization in the ovary undergoes sequential and significant changes, suggesting distinct functional roles of this transcription factor during oocyte maturation.

### 3.4. Cic is involved in the oocyte development of *B. germanica*

To unveil the functions of Cic in the panoistic ovaries of *B. germanica*, 1µg of ds*cic* was injected into two different experimental models: 0-day-old adult (ds*cic*-treated n = 45; dsMock n = 20) and 0-day-old sixth instar nymphs (ds*cic*-treated n = 65; dsMock n = 20). In ds*cic*-treated adult females just after the emergence, the ovarian mRNA levels for *cic* were measured 48h after the treatment, exhibiting 27% reduction. This depletion was sufficient to perturb oviposition and embryogenesis in the early stages. From the 45 treated females, 18% did not oviposit and 4% died before oviposition. Those treated females that oviposited (78%, n = 35), 60% dropped the ootheca 48 – 72 hours after its formation. The remaining females maintained the ootheca attached to the genital pouch, which gave rise to nymphs that emerged normally without delays.

The females treated as 0-day-old sixth instar nymphs (n = 65), molted normally to the adult 8 days later, similar to the dsMock-treated nymphs. At the end of the gonadotrophic cycle, all dsMock-treated females oviposited and had a normal ootheca, whereas none of the ds*cic*-treated females did. Ovarian *cic* mRNA levels were measured every 48 h during the nymphal instar and just after the adult emergence (Fig. 4A). The results indicated that in 8-day-old sixth instar nymphs treated with ds*cic*, the expression of *cic* mRNA tended to decrease compared to those treated with dsMock (Fig. 4A). In addition, in these 8-day-old ds*cic*-treated nymphs (Fig. 4B), no Cic protein labeling was detected in the BOF. Only a faint signal was observed at the basal pole of the follicular cells (Fig. 4B and Fig. S2). Conversely, in 8-day-old dsMock-treated nymphs, Cic labeling appeared in the BOF ooplasm and the basal pole of the follicular cells (Fig. 4C, and Fig. S2).

**Figure 4.**
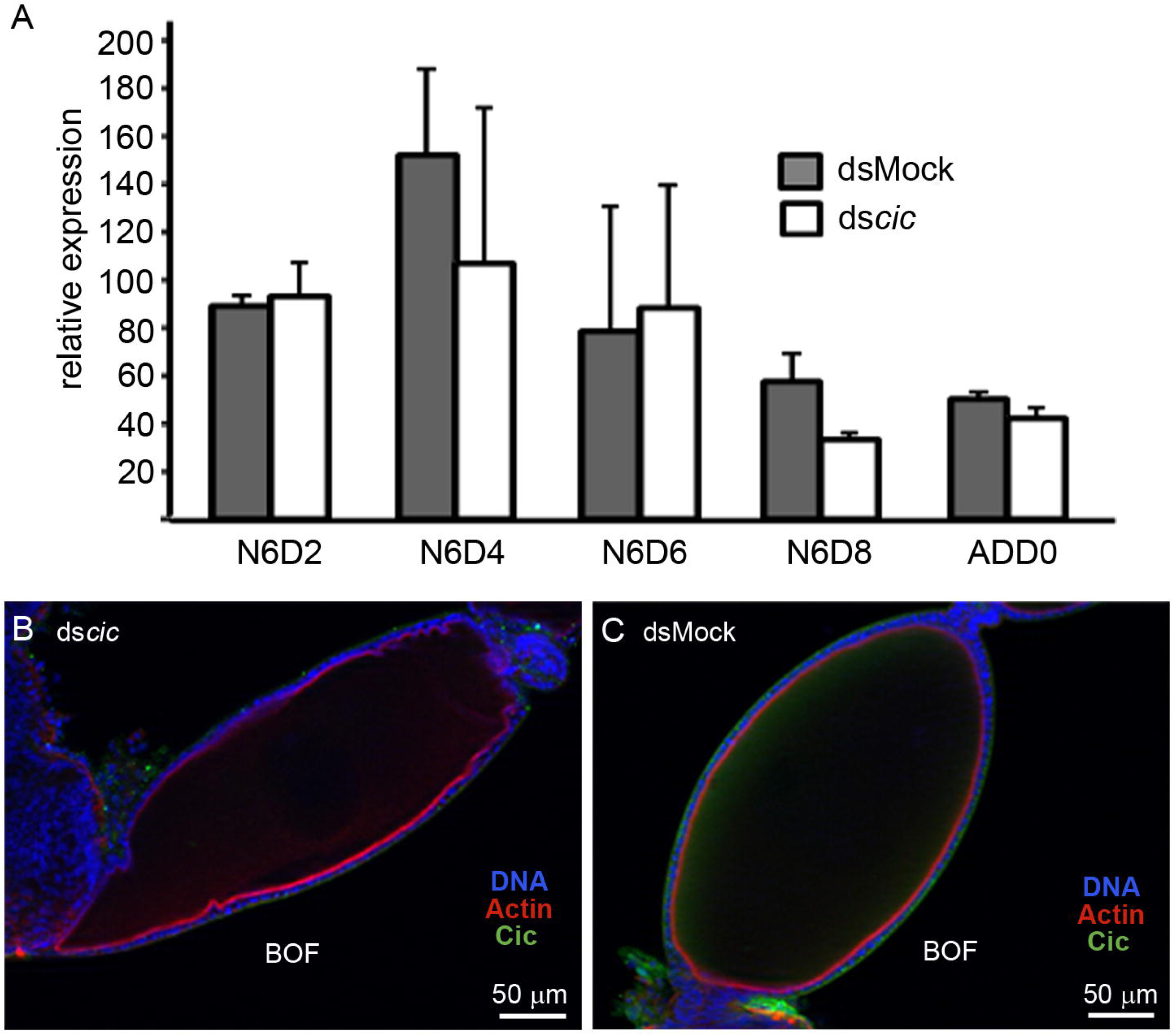
**Capicua (Cic) in ovaries of ds*cic-*treated females**. (A) Relative expression of *cic* mRNA in ovaries of sixth instar nymphs and newly emerged adult ds*cic*-treated. ds*cic* was applied to nymphs just after the molt to last instar and the RNA levels were measured at different developmental stages (N6D2, N6D4, N6D6, N6D8 and AdD0, n = 3 for each age and treatment). Data are expressed as copies of *cic* per 1000 copies of *actin-5c*. (B) Immunolocalization of Cic protein in basal follicles of 8-day-old ds*cic*-treated (n = 3) and (C) in dsMock-treated sixth instar nymph (n = 3). The anterior pole of the basal follicle in A and B is towards the right. Cic: green, F-actin: red, DAPI: blue. In Figure S2 the labeling intensity for each image is show.

The most evident effect observed in these ds*cic*-treated females was the change of the BOF shape, which appeared elongated with defects affecting mainly the anterior and posterior poles (Fig. 4B) a morphology that is very different from that of the dsMock-treated females, which have the typical elliptical shape (Fig. 4C). A new batch of 0-day-old ds*cic*-treated (n = 20) and dsMock-treated (n = 20), sixth instar nymphs, was prepared to observe the ovarian follicles at different ages, in the last instar nymphs and adults.

In 8-day-old ds*cic*-treated sixth instar nymphs, the BOF appeared elongated (Fig. 4B) compared to dsMock-treated nymphs (Fig. 4C). The anterior pole of the oocyte became narrow extending into the lumen of the stalk (Fig. 5A and B, arrowheads) whereas, the posterior pole showed a “fishtail-like” morphology (Fig. 5C and D, arrowheads). After the molt, in 3-day-old adults, the BOF of ds*cic*-treated females appeared even more elongated (Fig. 5E) and the apical pole of the oocyte that was entering the lumen of the stalk, appeared engulfed by the follicular cells located in the anterior pole of the ovarian follicle (Fig. 5F and G, arrowheads). Later, in 7-day-old adults, it is observed that the ovarioles of females treated with ds*cic* have lost their basal and subbasal oocytes, leaving only the sheath of the follicular epithelium that surrounded them (Fig. 5H, arrowheads). Additionally, the remaining distal oocytes in the vitellarium, display an unusual hourglass shape, probably due to the generalized enrichment of F-actin in the oocyte membrane and ooplasm (Fig. 5H arrow, and I).

**Figure 5.**
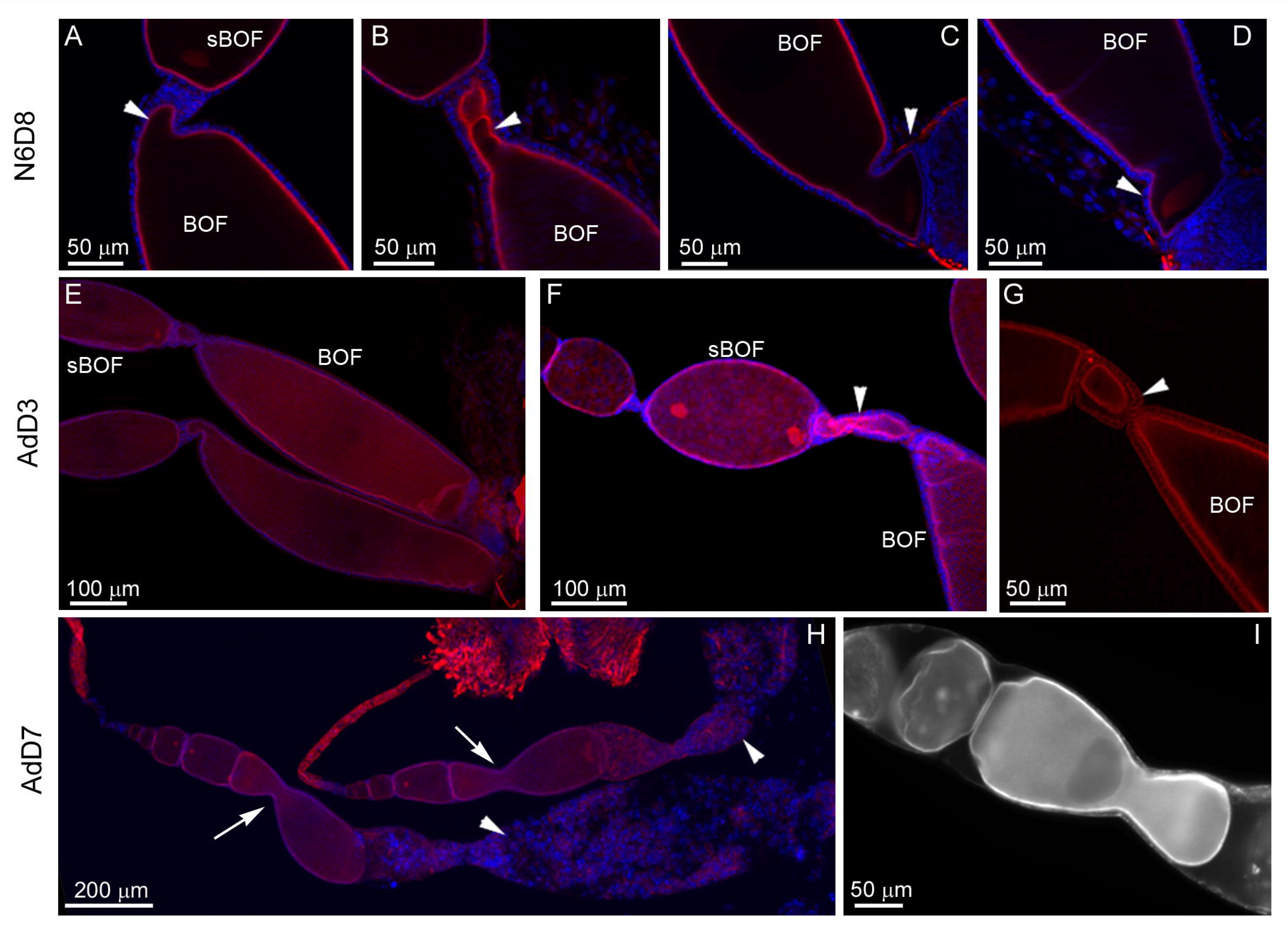
Depletion of *cic* in basal ovarian follicles of different ages. Last instar nymphs were treated the day of emergence, and the phenotypes analyzed at different ages. (A, B) Basal ovarian follicles (BOF) from 8-day-old ds*cic*-treated sixth instar nymphs, showing different degrees of narrowing in the anterior pole (arrowheads). (C, D) BOFs exhibiting the “fishtail-like” phenotype, with a concentration of F-actin in the posterior pole (arrowheads). (E-G) Ovarioles from 3-day-old ds*cic*-treated adults, showing the elongated shape of the BOF (E), and the engulfment of the anterior pole of the basal oocyte by the follicular epithelia (F, G; arrowheads). (H) Ovarioles from a 7-day-old ds*cic*-treated adult. The basal and sub-basal oocytes were prematurely oviposited (arrowheads) before completing their development, leaving the follicular epithelia that surround those oocytes. In these ovarioles, the new first oocyte shows an hourglass shape (arrows). (I) detail of the new first oocyte showing an hourglass shape. In addition, it was not possible to detect the stalk separating the ovarian follicles. The anterior pole is towards the left. F-actin: red (except in I, which is white), DAPI: blue.

Usually, in last instar nymphs, at both poles of the younger oocytes in the vitellarium, is observed an accumulation of F-actin, constituting the microtubule organization center (MTOC) (Fig. 6A, arrowheads), with a network of F-actins connecting both MTOCs (Fig. 6A), which usually disappears in the basal oocytes. However, in the basal oocyte of 8-day-old ds*cic*-treated nymphs, the MTOC remains present and strongly labeled (Fig. 6B and C). It is also preserved and significantly larger in the BOF of 3-day-old adults from the ds*cic*-treated group (Fig. 6D). The presence of the MTOCs at non-corresponding ages may be the cause of changes in the shape of the ovarian follicles.

**Figure 6.**
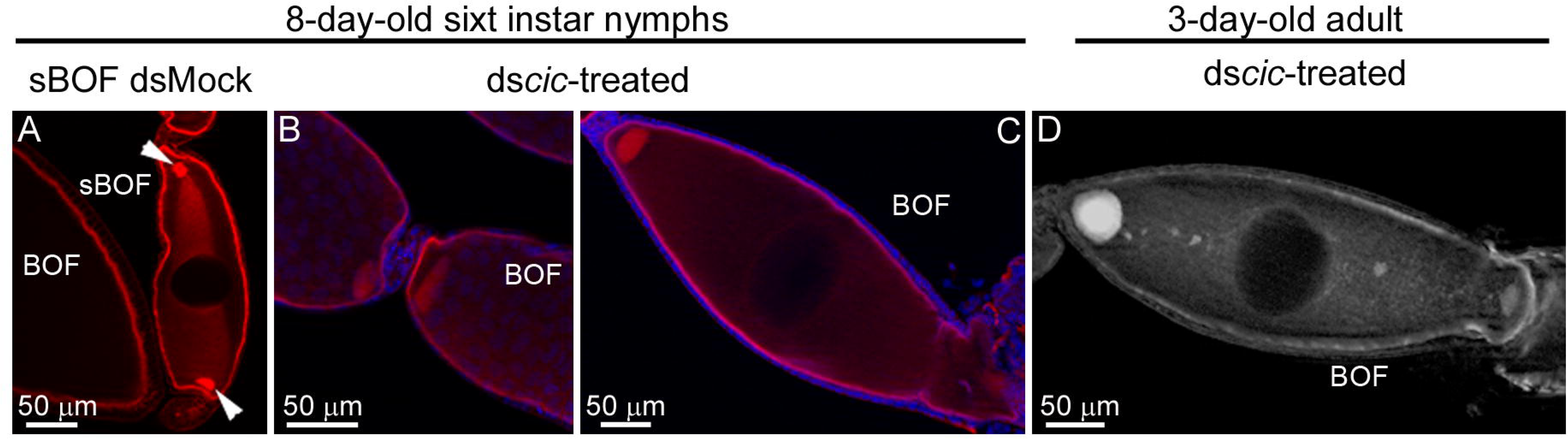
**MTOC in ovarian follicles from N6D8 and AdD3 treated with ds*cic* newly-emerged sixth instar nymph.** (A) Sub-basal ovarian follicle from an 8-day-old-dsMock treated sixth instar nymphs, showing the MTOCs placed in the anterior and posterior oocyte poles (arrowheads), interconnected by and F-actin network. (B, C) BOF from 8-day-old ds*cic*-treated sixth instar nymph, showing an abnormally enlarged MTOC in the anterior pole of the oocytes. (D) BOF from a 3-day-old ds*cic*-treated adult showing the network of F-actin, with a big MTOC in the anterior pole of the oocyte, when, usually at this age, the MTOC and the F-actins network, are not detected. The anterior pole is towards the left. F-actin: red (except in D that is white), DAPI: blue.

### 3.5. Depletion of *cic* mRNA affects the immature ovarian follicles

In the young ovarian follicles from 8-day-old ds*cic*-treated sixth instar nymphs (Fig. 7), the nucleus in these oocytes is not in a central position, as occurs in controls (Fig. 7A). Rather, it takes a posterior position, in the oocytes within the ovariole (Fig. 7B-D, arrowheads). It seems that the nucleus was pushing the oocyte towards the posterior pole of the ovarian follicle, and as a consequence, many oocytes can be encapsulated in one ovarian follicle (Fig. 7D). This is facilitated by the absence of follicular cells in the poles and the lack of stalk cells separating young ovarian follicles, which determined the fusion of the oocyte membranes, thus giving the aspect of a single oocyte with more than one nucleus (Fig. 7B-D, arrowheads). Moreover, the absence of the stalk connecting successive ovarian follicles leads to a general disorganization of the ovariole, especially affecting recently formed ovarian follicles, that are ready to leave the germarium. As a consequence of these dysfunctions, in older adult females (7-day-old), the malformed BOF has fallen, and is lost into the oviduct, before completing maturation (Fig. 5H).

**Figure 7.**
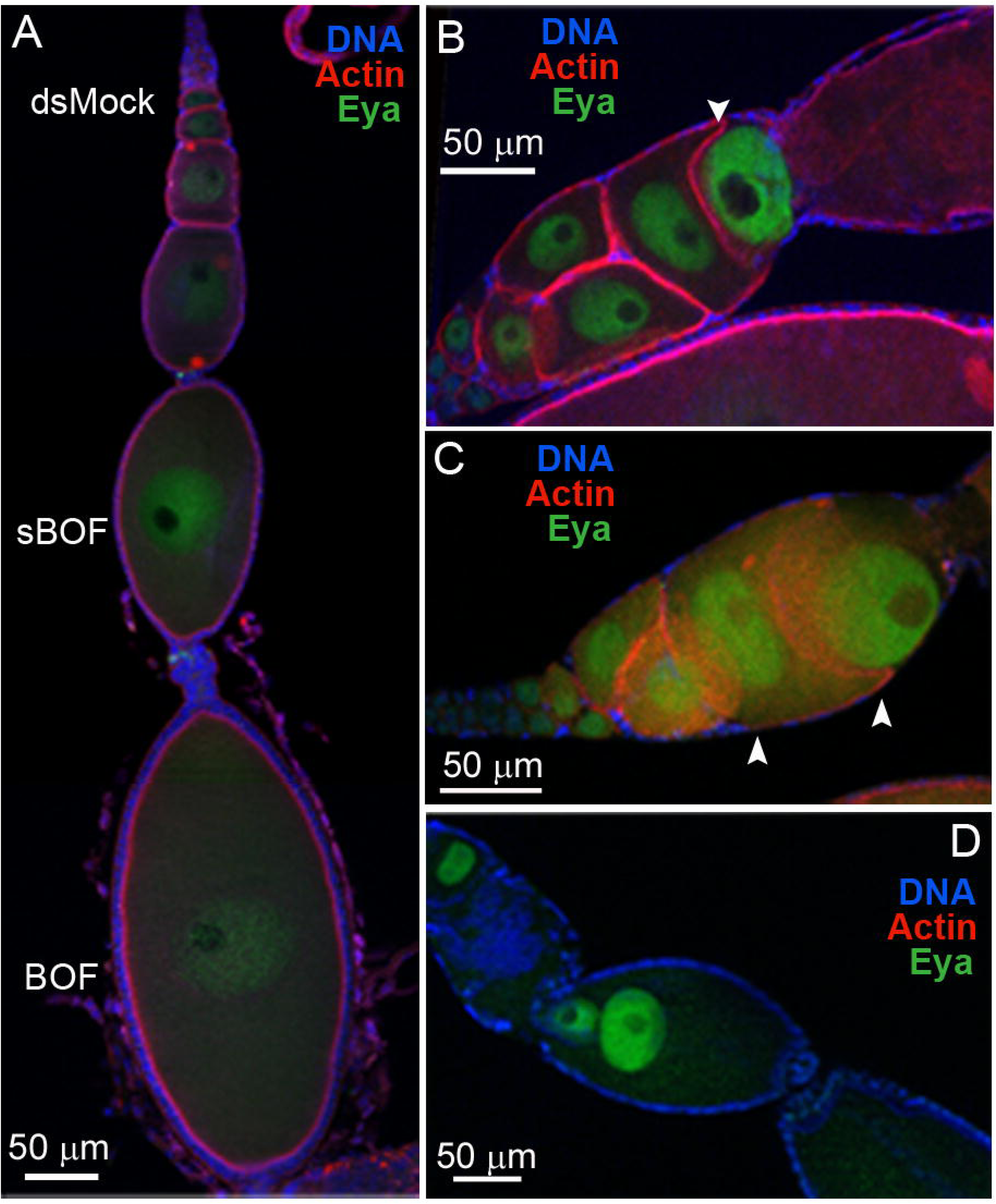
Localization of the oocyte nucleus in ds*cic*-treated sixth instar nymphs. (A) Ovariole from an 8-day-old dsMock-treated nymph, showing the normal localization of oocyte nucleus in the different ovarian follicles along the ovarioles (BOF: basal ovarian follicle; sBOF: sub-basal ovarian follicle). (B, C) Disorganization of the ovarian follicles in the vitellarium due to the stalk loss showing the mislocalization of the oocyte nucleus (The arrowhead indicates the expected position of the stalk, which is absent). (D) Fusion of two oocytes resulting in a single ovarian follicle containing two nuclei. In B the anterior pole is up towards the left. The oocyte nucleus is labeled by anti-Eya antibody (green); F-actin: red; DAPI: Blue.

### 3.6. Cic interacts with other signaling pathways in the ovary

The complex phenotypes produced by the depletion of *cic* mRNA suggests that Cic protein could interact with other pathways related to cell proliferation and stalk formation. As Cic is a downstream effector of EGFR signaling, we analyzed the EGFR in 8-day-old sixth instar ds*cic*-treated nymphs. The expression of *EGFR* in ovaries tended to decrease, although the differences compared with the control group were not statistically significant (Fig. 8A). However, the localization of EGFR protein in ovaries of ds*cic*-treated nymphs, appeared modified compared to dsMock-treated nymphs (Fig. 8B-D). In ds*cic*-treated females, EGFR labeling was more restricted to the mid-ventral region of the basal oocyte and around the nucleus (Fig. 8C-D). In contrast, in dsMock-treated nymphs, EGFR localization is less restricted and spreads over the cytoplasm and within the nucleus, with a higher concentration close to the nucleus membrane (Fig. 8B).

**Figure 8.**
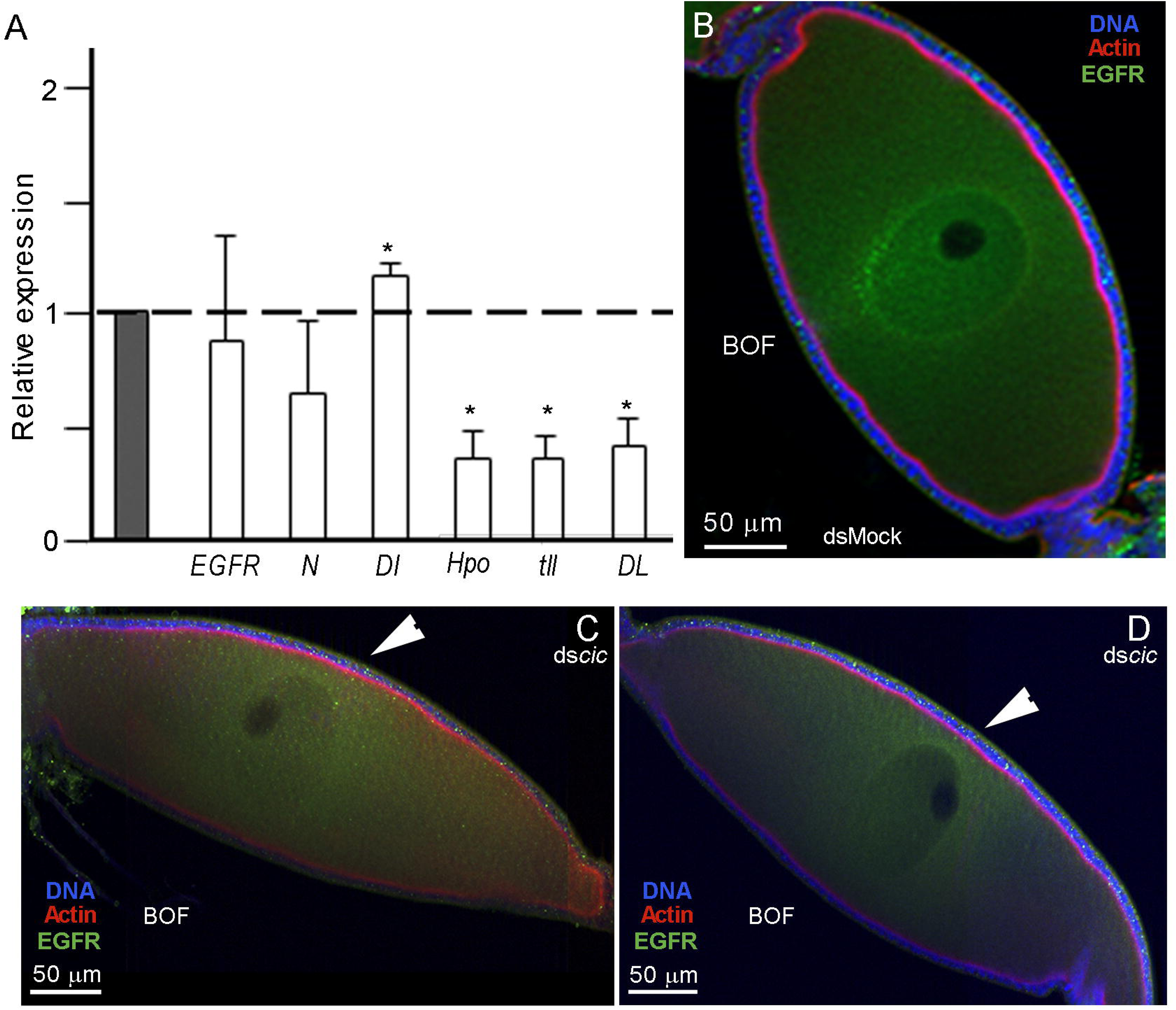
Genes involved in oocyte growth and development. (A) Relative expression of *EGFR, notch* (*N*)*, delta* (*Dl*)*, hippo* (*hpo*)*, tailless* (*tll*), and *dorsal* (*DL*) in ovaries from 8-day-old ds*cic*-treated sixth instar nymphs. Data represent normalized values against dsMock controls (reference value = 1) (n = 3) (*) indicates statistically significant change P(H1) = 0.0001. (B) Basal ovarian follicle from 8-day-old dsMock-treated sixth instar nymph, showing the EGFR protein with high labeling on nuclear membrane and inside the oocyte nucleus. (C-D) Basal ovarian follicles of a ds*cic*-treated nymph showing changes in EGFR localization. In these follicles EGFR labeling appeared restricted to the central region of the oocyte around the nucleus and towards the oocyte ventral side (ar-rowhead). F-actin: red; DAPI: Blue, EGFR protein: Green.

The absence of stalk between ovarian follicles in ds*cic*-treated specimens, and the reduced number of follicular cells in some areas of the ovarian follicles, suggest us to measure the expression of some components of the Hippo and Notch pathways. In *cic* knockdown nymphs, *hippo* (*hpo*) was significantly down-regulated (0.351-fold change) (Fig. 8A). We also measured the expression of *Notch* (*N*) and *Delta* (*Dl*), from the Notch pathway (Irles et al., 2016; Irles & Piulachs, 2014), and as expected, the expression of *N* tended to decrease, while its ligand *Dl* was significantly up-regulated (1.161-fold change; Fig. 8A), suggesting that the Hippo and Notch pathways respond to Cic signaling.

Finally, the modifications observed in the anterior-posterior poles in ovarian follicles from ds*cic*-treated sixth instar nymphs suggested looking for genes that drive the terminal position in *D. melanogaster* egg chambers and embryos. Therefore, we examined the response of *tailless* (*tll*) and *Dorsal* (DL) (Fig. 8A) in ds*cic*-treated nymphs. The expression levels of *tll* were significantly down-regulated (0.355-fold change). While the expression of the transcription factor DL, which is required for the determination of dorsal-ventral polarity, decreased significantly (0.408-fold change) (Fig. 8A), reinforcing the role of *cic* in the panoistic ovary, maintaining the polarity of the ovarian follicles.

## 4. Discussion

Cic is a transcription factor that is highly conserved among animals, from cnidarians to vertebrates (Lam et al., 2006; C. J. Lee et al., 2002). The function of Cic has been extensively studied in *D. melanogaster*, where two isoforms have been identified (Jiménez et al., 2012; Rodrıguez-Muñoz et al., 2022). One is the Cic-S, considered the canonical version, which appears mainly in dipterans (Fig. S1). The longer isoform, found in most insect species, including *B. germanica*, was described in *D. melanogaster* as responsible for maintaining the correct somatic and germinal cell cycle in the ovary (Rodrıguez-Muñoz et al., 2022). These insect Cic sequences contain the typical domains for the long Cic isoform (Forés et al., 2015), including the N2 motif, described previously as Diptera exclusive (Forés et al., 2015). The phylogenetic analysis of the insect Cic sequences suggests that the long isoforms could be ancestral, and the Cic-S was an innovation in some insect groups.

The *B. germanica cic* is expressed in the brain and ovaries, similar to the *D. melanogaster* Cic-L isoform, affecting brain size and development (Yang et al., 2016). The high levels of *cic* expression in the *B. germanica* brain suggest that Cic may participate in developing the central nervous system. However, we focused our research on the expression and function of Cic in the panoistic ovary of *B. germanica*.

The role of Cic in insect oogenesis and embryo development has been thoroughly documented in *D. melanogaster*. In this species, Cic participates in the specification of terminal regions within the embryo; the loss of Cic function results in dorsalization of both the eggshell and the embryo (Atkey et al., 2006; Goff et al., 2001; Jiménez et al., 2000; Rodríguez-Muñoz et al., 2022), affecting the anterior-posterior embryonic axis (Andreu et al., 2012). In *T. castaneum*, a species with meroistic telotrophic ovaries and a short germband embryo, Cic is essential for anterior-posterior patterning without any significant impact on dorsoventral pattern formation, and its depletion causes the terminal anlagen to enlarge (Pridöhl et al., 2017). In the panoistic ovary of *B. germanica*, Cic protein accumulates in the basal oocytes of sixth-instar nymphs as they progress in the instar. However, after the imaginal molt, Cic labeling in the basal oocytes becomes weak, suggesting that, in the panoistic ovary, the most important steps for oogenesis and further oocyte maturation occur during the last nymphal instar. Thus, when the female reaches the adult stage, the basal follicles are ready to incorporate storage proteins into the oocyte and to complete growth (Belles et al., 2024; Ciudad et al., 2006; Herraiz et al., 2014; Irles and Piulachs, 2014; Tanaka and Piulachs, 2012). According to our findings, Cic plays a crucial role in oogenesis, which occurs before the ovarian follicle reaches maturity, and its depletion causes female sterility. Equivalent observations have been reported for *D. melanogaster* (Jiménez et al., 2000), where Cic has a fundamental function in oocyte development, determining female sterility when depleted. However, the mechanisms underlying this macroscopic phenotype differ between the two species with two ovary types.

*D. melanogaster* Cic mutants can oviposit, although eggs have defects in the eggshell, and the embryos begin to develop. However, they do not complete the embryogenesis and cannot hatch (Goff et al., 2001; Rodríguez-Muñoz et al., 2022). Conversely, in *B. germanica*, *cic*-depleted females do not lay eggs correctly because the oocyte cannot complete its development. In *B. germanica*, Cic is required to maintain the anterior-posterior axis since depletion of *cic* mRNA triggers morphological modifications in the anterior-posterior poles of the BOFs, resulting in significant changes in the distribution of F-actin microfilaments, in both the oocytes and the follicular cells.

Actin fibers are essential components of the cytoskeleton and play a crucial role in various cellular processes, such as cell division, cell shape maintenance, oocyte growth, and establishing oocyte polarity (Riparbelli & Callaini, 1995; Sun & Schatten, 2006). In last instar nymphs, the MTOCs in the growing oocytes are located at the anterior and posterior poles. These MTOCs disappear from the basal oocytes in the days leading up to the molt into adulthood and are never detected in adult oocytes. However, in females treated with ds*cic*, the MTOCs remain at the poles of the basal oocyte during the transition from nymph to adult. The role of the MTOC in organizing the microtubule network is crucial for the distribution of mRNAs and proteins within the oocyte, as shown in studies involving *D. melanogaster* (Cooley & Theurkauf, 1994; Roth & Lynch, 2012). Although a network of F-actin can be observed connecting the two MTOCs at the poles in BOFs from adult ds-*cic*-treated, we cannot provide information regarding the function of MTOCs in the distribution of proteins or mRNAs in panoistic ovaries. The varying prevalence of MTOCs at different stages suggests that this fact may influence the shape changes observed in ovarian follicles.

Cic is a downstream effector of EGFR signaling that controls cell growth, differentiation, and survival in different contexts (Goff et al., 2001; Elshaer and Piulachs, 2015). In *B. germanica* EGFR-depleted nymphs, the anterior-posterior polarity is disrupted, leading to ovarian follicles, and ovariole disorganization (Elshaer & Piulachs, 2015). The restriction of EGFR protein to the mid-ventral region might explain the fusion of immature oocyte membranes in ds*cic* specimens of *B. germanica*, determining the fusion of some oocytes and suggesting that Cic in the panoistic ovary of *B. germanica* is necessary to transduce the EGFR signaling, as happens in *D. melanogaster*, where Cic is required to translate the EGFR signaling to pipe and establishing the nuclear dorsal gradient (Moussian & Roth, 2005). In addition, the relationship between EGFR signaling and cell proliferation has been established in different species. In *D. melanogaster*, the inhibition of Cic by EGFR leads to the de-repression of hpo and subsequently influences cell proliferation (Herranz et al., 2012). Conversely, our findings in *B. germanica* indicate that the depletion of *cic* results in the down-regulation of *hpo*. However, the follicular epithelium continues to proliferate, likely because Notch signaling within the ovarian follicle sustains an adequate level of proliferation, as previously reported (Irles & Piulachs, 2014). This indicates that the follicular epithelium can continue to proliferate even with low Hpo levels, due to the ongoing activity of N in the follicular cells. Previously, we demonstrated that Dl in young females is accumulated at the anterior and posterior poles of basal ovarian oocytes (Irles et al., 2016), interacting with N to determine specific cell populations in de BOF. The depletion of Dl in ds*cic*-treated females could also be responsible for the phenotypes observed in the BOFs.

Depletion of Cic in *B. germanica* seems to modify the expression of those genes that in *D. melanogaster* are required to establish the anterior-posterior polarity or the DV axis (like tailless and dorsal) (Lynch & Roth, 2011; Roth & Lynch, 2009), as they were down-regulated in the cockroach. It will be necessary to investigate the function of all these genes in a panoistic ovary, where the polarity of the oocyte differs greatly from that of a meroistic oocyte, and unveil how its function has been modified through evolution.

## Acknowledgments

The work was funded with the grant PID2021-122316OB-I00 from the MICIU/AEI /10.13039/501100011033 and ERDF, a way of making Europe. We are grateful to Xavier Belles for the critical reading of the manuscript.

Declarations of interest: none

---

**Figure S1.**
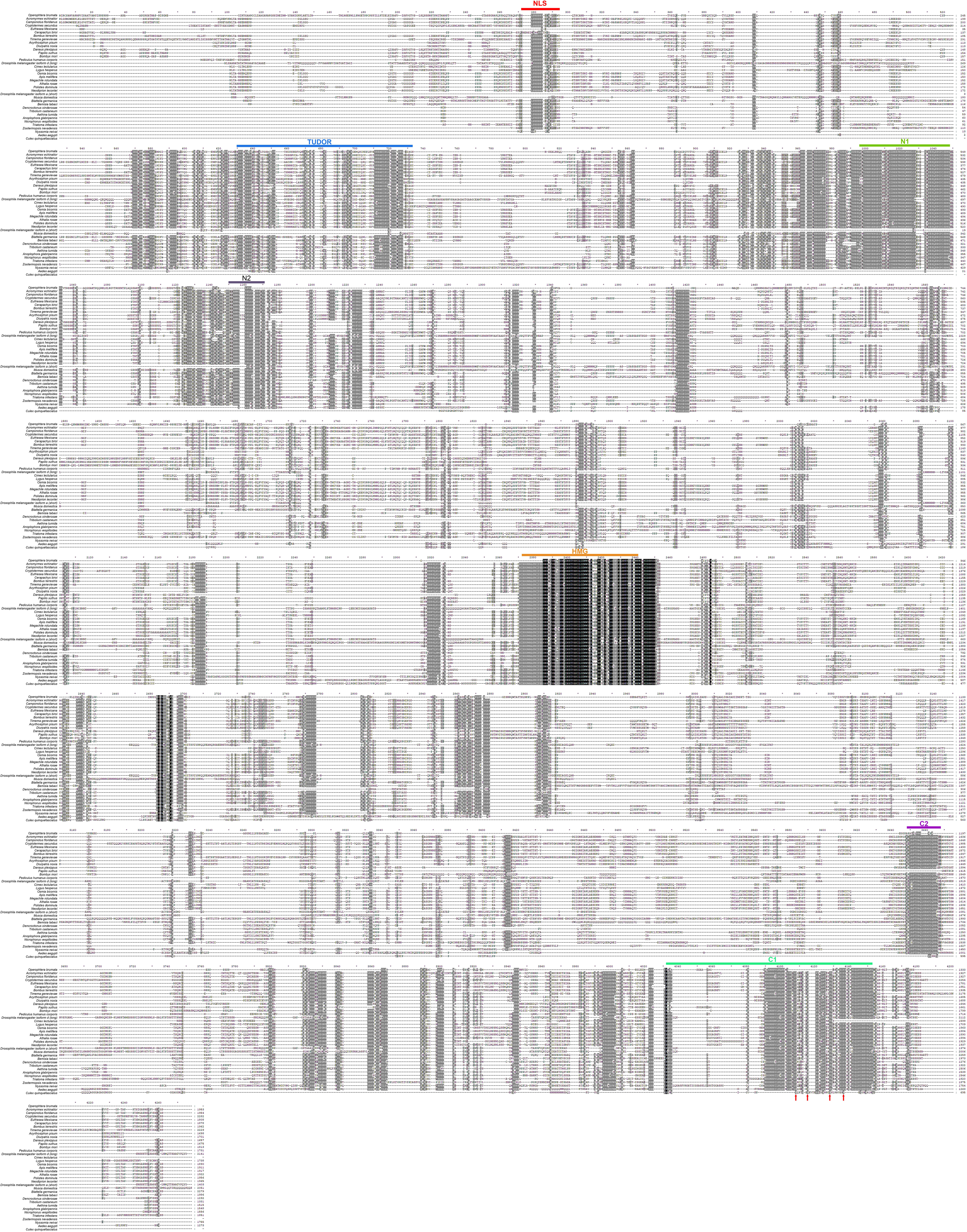
Multiple Sequence alignment of the Capicua protein across different insect species, using MAFFT (Katoh et al., 2013). Key functional domains—NLS, Tudor, N1, N2, HMG, C1, and C2—are highlighted in distinct colors (red, blue, green, violet, orange, purple, and fluorescent green, respectively) to indicate their locations and relative conservation. Highly conserved regions are marked, emphasizing the evolutionary stability of these functional motifs. The characteristic phenylalanine residues with the C1 domain are specifically indicated (red arrows). Accession numbers from different species are provided in Supplementary Table 1

**Figure S2.**
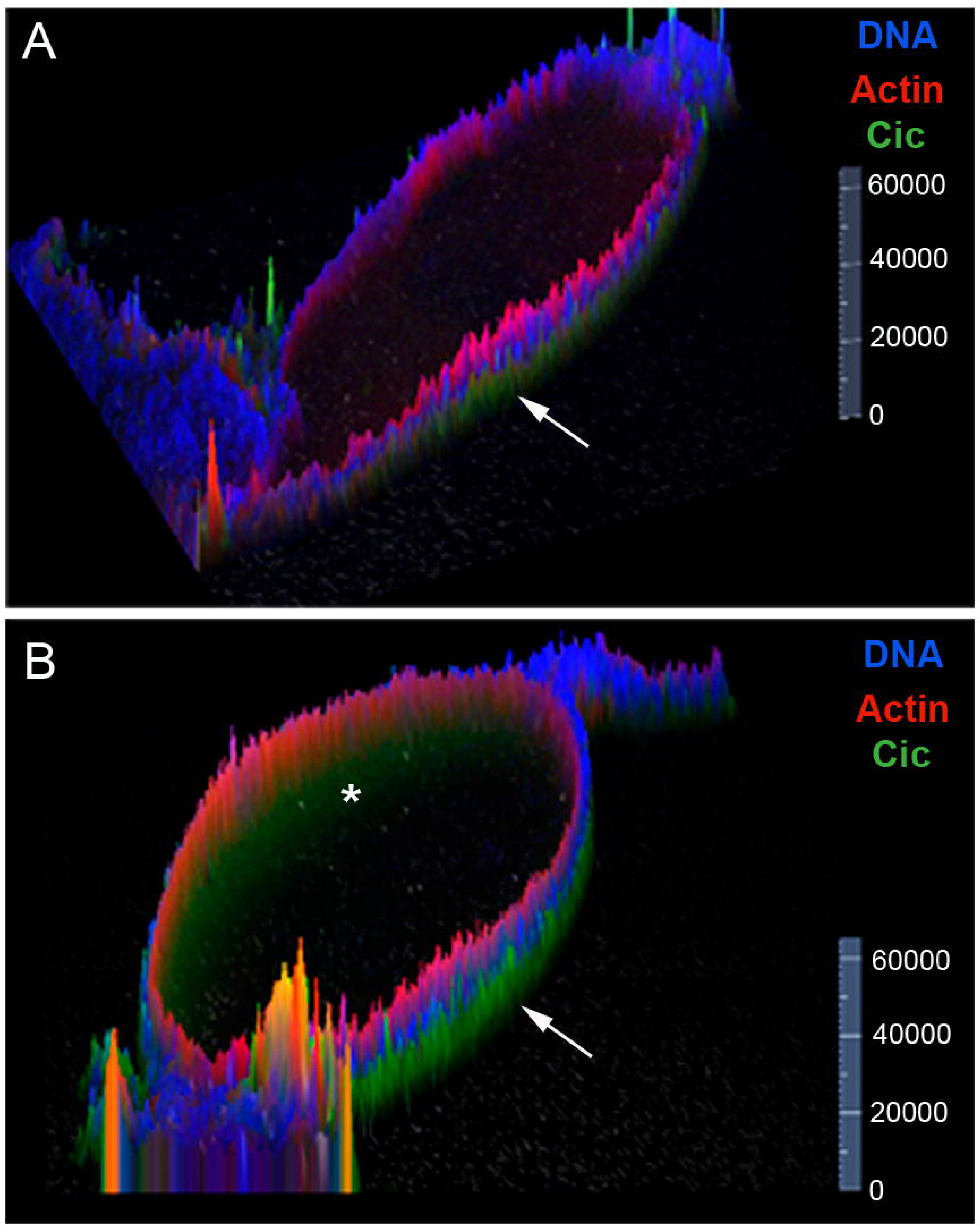
Cic protein in basal ovarian follicles of dscic treated females. (A) Cic protein in basal follicles of 8-day-old dscic-treated and (B) in dsMock-treated sixth instar nymph (Images from Figure 4). The relative fluorescence intensities of the different fluorophores in the Figure 4B and C,are shown in B’ and C’. It was graphically quantified using the Zen 2.5D analysis tool (Carl Zeiss Microscopy).The anterior pole of the basal follicle in A and B is towards the right. Cic: green, F-actin: red, DAPI: blue.

