## Supplementary data for "The transcription factor Capicua maintains the oocyte polarity in the panoistic ovary of the German cockroach"

| **Order** | **Species** | **Accession number** |
| --- | --- | --- |
| *Blattodea* | *Blattella germanica* | LN623700 |
| *Blattodea* | *Cryptotermes secundus* | XP_023709425.1 |
| *Blattodea* | *Zootermopsis nevadensis* | KDR16881.1 |
| *Coleoptera* | *Tribolium castaneum* | XP_968497.2 |
| *Coleoptera* | *Dendroctonus ponderosae* | ERL94807.1 |
| *Coleoptera* | *Anoplophora glabripennis* | XP_018578847.1 |
| *Coleoptera* | *Nicrophorus vespilloides* | XP_017787088.1 |
| *Coleoptera* | *Aethina tumida* | XP_019866354.1 |
| *Diptera* | *Drosophila melanogaster isoform a (short)* | NP_524992.1 |
| *Diptera* | *Drosophila melanogaster isoform d (long)* | NP_001247203.1 |
| *Diptera* | *Culex quinquefasciatus* | XP_001862155.1 |
| *Diptera* | *Aedes aegypti* | XP_001649010.1 |
| *Diptera* | *Musca domestica* | XP_011294382.2 |
| *Diptera* | *Nyssomyia neivai* | JAV06247.1 |
| *Hemiptera* | *Diurpahis noxia* | XP_015374761.1 |
| *Hemiptera* | *Triatoma infestans* | JAC14751.1 |
| *Hemiptera* | *Besimia tabaci* | XP_018912315.1 |
| *Hemiptera* | *Acyrthosiphon pisum* | XP_003241821.1 |
| *Hemiptera* | *Lygus hesperus* | JAQ09670.1 |
| *Hemiptera* | *Cimex lectularius* | XP_014242481.1 |
| *Hymenoptera* | *Acromyrmex echinatior* | EGI66064.1 |
| *Hymenoptera* | *Camponotus floridanus* | EFN68385.1 |
| *Hymenoptera* | *Bombus terrestris* | XP_003393598.1 |
| *Hymenoptera* | *Apis mellifera* | XP_003249448.1 |
| *Hymenoptera* | *Megachile rotundata* | XP_003706301.1 |
| *Hymenoptera* | *Osmia bicornis* | XP_029031934.2 |
| *Hymenoptera* | *Polistes dominula* | XP_015187054.1 |
| *Hymenoptera* | *Athalia rosae* | XP_012260851.1 |
| *Hymenoptera* | *Neodiprion lecontei* | XP_015521021.1 |
| *Hymenoptera* | *Eufriesea Mexicana* | OAD54499.1 |
| *Hymenoptera* | *Cerapachys biroi* | XP_011333295.1 |
| *Lepidoptera* | *Danaus plexippus* | EHJ68194.1 |
| *Lepidoptera* | *Bombyx mori* | XP_004931421.1 |
| *Lepidoptera* | *Papilio xuthus* | KPI94261.1 |
| *Lepidoptera* | *Operophtera brumata* | KOB70364.1 |
| *Phasmotodea* | *Timema genevievae* | CAD7586469.1 |
| *Psocodea* | *Pediculus humanus corporis* | XP_002423691.1 |

**Supplementary table 1**. Name of the species used in the phylogenetic analysis, with corresponding accession number for Cic sequence. The sequence was obtained from NCBI database.

**Supplementary table 2**: **Primer sequence used for qRT-PCR and RNAi experiments.** The accession numbers of studied sequences are indicated. F: Primer forward. R: Primer reverse. RT: primer used in real-time PCR; dsRNA: primers used to prepare the dsRNA.

|  | **Accession number** | **Primer name** |  | **Primer sequence** | **Amplicon length (bp)** |
| --- | --- | --- | --- | --- | --- |
| 1 | LN623700 | *cic*-RT | F  R | 5’ AACCCGCAAGGTTGTCAGT 3’  5’ ACTCTGCTGCTCAAGCACAA 3’ | 150 |
| 2 | LN623700 | *cic*-dsRNA1 | F  R | 5’ ACTTGAGAGGAGAACCAGA 3’  5’ ATGGACAGGGTTCTCGAAAC 3’ | 324 |
| 3 | LN623700 | *cic* -dsRNA2 | F  R | 5’ CCAGCAAGGTTAACATCTTGCAA 3’  5’ CAACTCGACCGAAGTCAACAG 3’ | 392 |
| 4 | LN623701 | *EGFR*-RT | F  R | 5’ GAGTACAAAGCAGCAGGAG 3’  5’ CCAACATCGGATATTCACTCAC 3’ | 78 |
| 5 | LN623703 | *DL*-RT | F  R | 5’ GGTTTCTCTCATCGCAGTCA 3’  5’ CAGTGGTGTCTGAACCCATC 3’ | 132 |
| 6 | LN623702 | *Tll*-RT | F  R | 5’ GACAGCGTCAGTACGTTTGC 3’  5’ ATGAACAAGGATGCGGTACA 3’ | 130 |
| 7 | HF969255 | *N*-RT | F  R | 5’-GCTAAGAGGCTGTTGGATGC-3’  5’-TGCCAGTGTTGTCCTGAGAG-3’ | 55 |
| 8 | HF969256 | *Dl*-RT | F  R | 5’-CCACTACAAGTGTTCGCCAA-3’  5’-TACCTCTCGCATTCGTCACA-3’ | 180 |
| 9 | HF969251 | *Hpo*-RT | F  R | 5’-GACATTTGGAGCCTTGGCAT-3’  5’-AGGTTTCCCTTCAGCCATTTC-3’ | 51 |
| 10 | AJ862721 | *actin-5c*-RT | F  R | 5’-AGCTTCCTGATGGTCAGGTGA-3’  5’-ACCATGTACCCTGGAATTGCCGACA-3’ | 213 |
